# Neutral sphingomyelinase 1 regulates cellular fitness at the level of ER stress and cell cycle

**DOI:** 10.1101/2022.02.23.481585

**Authors:** Dolma Choezom, Julia Christina Gross

**Affiliations:** Developmental Biochemistry, University Medical Center Goettingen, Goettingen, Germany; Hematology and Oncology, University Medical Center Goettingen, Goettingen, Germany; Health and Medical University, Potsdam, Germany

**Keywords:** Cell cycle arrest, Wnt-signaling, Ceramide, PI3K/AKT, ER stress, LAMP1, translation

## Abstract

Neutral sphingomyelinase 1 (nSMase1) belongs to the sphingomyelinase enzyme family that hydrolyzes sphingomyelin to produce signaling active lipid ceramide and phosphorylcholine. The molecular characterization and biological function of nSMase1 remain poorly studied. Here, we report that nSMase1 (gene name: SMPD2) knockdown reduces LAMP1 at the mRNA levels and is required for initiating a full-potential unfolded protein response under ER stress. Additionally, SMPD2 KD dramatically reduces the global protein translation rate. We further show that SMPD2 KD cells are arrested in the G1 phase of the cell cycle and that two important cell cycle regulating processes - PI3K/Akt pathway and Wnt signaling pathway are altered. Taken together, we propose a role for nSMase1 in buffering ER stress and modulating cellular fitness via cell cycle regulation.

## Introduction

Sphingomyelin, a major lipid constituent of the plasma membrane, through its catabolic metabolism generates several bioactive signaling lipids: ceramide, sphingosine, and sphingosine1-phosphate (Hannun & Obeid, 2008). Sphingomyelindiesterases (SMPDs) or sphingomyelinases (SMases) are a family of enzymes that hydrolyze the phosphodiester bond of the sphingomyelin to produce ceramide and phosphorylcholine (Airola & Hannun, 2013). The sphingomyelinases are broadly divided into three major classes: acid sphingomyelinases, alkaline sphingomyelinases, and neutral sphingomyelinases based on their pH optima (Airola & Hannun, 2013). In mammals, four neutral sphingomyelinase members have been identified, nSMase1 (Gene name: SMPD2) ((Tomiuk *et al*, 1998)), nSMase2 (SMPD3), nSMase3 (SMPD4), and mitochondrial associated nSMase (MA-nSMase; SMPD5) (Yabu *et al*, 2009; Wu *et al*, 2010).

We have previously shown that nSMase2 activity at the endosomal membrane regulates exosome secretion by counteracting V-ATPase-mediated endosomal acidification (Choezom & Gross, 2022). The number and different cellular localization of the neutral sphingomyelinase family members suggest distinct functions for each enzyme. NSMase1 shares identical domain architecture to ISC1, the yeast homolog to neutral sphingomyelinases (Tomiuk *et al*, 1998; Hofmann *et al*, 2000). In contrast to the overexpressed nSMase1 which localizes to the endoplasmic reticulum (Fensome *et al*, 2000; Rodrigues-Lima *et al*, 2000; Hofmann *et al*, 2000), endogenous nSMase1 exclusively localizes to the nuclear matrix (Mizutani *et al*, 2001). Although nSMase1 hydrolyzes sphingomyelin *in-vitro*, sphingomyelin metabolism remains unaffected upon its overexpression in cells (Tomiuk *et al*, 1998; Sawai *et al*, 1999). Likewise, nSMase1 mouse knockout showed no obvious phenotype with no detectable changes in sphingomyelin metabolism (Zumbansen & Stoffel, 2002). Instead of sphingomyelin, a study has shown that nSMase1 hydrolyzes lyso-platelet-activating factor (lyso-PAF) *in-vitro* and in cells (Sawai *et al*, 1999). This indicates the putative nSMase1 could be a lyso-PAF phospholipase C with lyso-PAF as its biological substrate (Sawai *et al*, 1999). Despite being the first neutral sphingomyelinase in mammals to get cloned and identified, the biochemical and molecular characterization of nSMase1 remains poorly studied.

In contrast to the above biochemical studies, several studies showed that ceramide generated by nSMase1 activity induces cellular death via different pathways under different stress conditions (Tonnetti *et al*, 1999; Jaffrézou *et al*, 1996; Jana & Pahan, 2004; Lee *et al*, 2004). JNK-signaling activates nSMase1 by phosphorylation to generate ceramide for apoptosis induction upon different environmental stresses such as heat shock and UV irradiation (Yabu *et al*, 2015). Given the insufficient and contrasting results on the role of nSMase1, we aimed to ascertain the biological role of nSMase1 by using siRNA-mediated gene knockdown (KD) in two human cell lines: HCT116, a human colorectal cancer cell line, and HeLa, a cervical cancer line.

Here, we discover that nSMase1 plays an important role in maintaining overall cellular homeostasis. SMPD2 KD reduces LAMP1 (lysosomal-associated membrane protein 1) proteins levels by downregulating its mRNA. And this dramatic reduction in LAMP1 protein has no apparent effect on the function of lysosomes as autophagy remained unaffected. Interestingly, SMPD2 KD cells are not only inefficient in activating a full-potential unfolded protein response (UPR) signaling upon ER stress but are arrested in the G1 phase of the cell cycle. We further show that two important cell cycle regulating processes - PI3K/Akt pathway and Wnt signaling pathway-are altered by SMPD2 KD. Specifically, SMPD2 KD affects PI3K/Akt signaling by significantly reducing the level of phosphorylated Akt. Most importantly, SMPD2 KD significantly reduces the Wnt signaling activity by reducing β-catenin protein levels in HeLa cells and Wnt3a in HCT116 at both protein and mRNA levels. Additionally, SMPD2 dramatically reduces the overall protein translation rate. These findings prove an important biological role for nSMase1 and provide strong foundations for further studies on dissecting its molecular function and pathway.

## Results

### SMPD2 Knockdown downregulates LAMP1 mRNA level

Sphinghomylinases are active at different cellular membranes (Airola & Hannun, 2013). We have previously shown that nSMase2 (gene name: SMPD3) activity at the endosomal membrane regulates exosome secretion by counteracting V-ATPase-mediated endosomal acidification (Choezom & Gross, 2022). Here, we analyzed the potential role of nSMase1 (gene name: SMPD2), a less well-understood member of the neutral sphingomyelinase family, in the regulation of endosomal trafficking decisions. SMPD2 knockdown (KD) significantly reduced HeLa cell viability when compared with control cells. In comparison, KD of SMPD3, NSMAF, an activating factor of nSMase2 (Philipp *et al*, 2010), and Syntenin (gene name: SDCBP), an adaptor protein involved in exosome biogenesis (Baietti *et al*, 2012), did not affect cell viability(Fig. S1A). These data indicate that SMPD2 is a vital gene for cellular survival. Upon SMPD3 KD, endosomal markers and acidified compartments accumulate intracellularly (Choezom & Gross, 2022). SMPD3 KD also slightly increased lysosomal protein LAMP1 levels, in contrast, SMPD2 KD strongly downregulated LAMP1 in both HeLa (Fig. 1A, Fig. S1B) and HCT116 (Fig. S1C). NSMAF and Syntenin (gene name: SDCBP) KD did not alter LAMP1 protein levels (Fig. 1A). Interestingly, SMPD2 KD did not affect lysosomal acidification as the total Lysotracker puncta per cell remained unaffected by SMPD2 KD (Fig. 1B, C). To directly determine whether the downregulation of LAMP1 protein upon SMPD2 KD affects lysosomal activity, we analyzed autophagy flux with two known markers – LC3B-II and P62 - by measuring their lysosomal turnover under both serum-fed and serum-starvation conditions with and without lysosomal inhibition with Bafilomycin-A (BAF) (Tanida *et al*, 2005). We found that P62 and LC3B-II flux remained unchanged upon SMPD2 KD under both basal and serum starvation conditions (Fig. 1D-F). These data allow two conclusions: Firstly, upon SMPD2 KD, downregulated LAMP1 does not affect lysosomal function. Secondly, as LAMP1 protein levels remained downregulated even upon lysosomal inhibition (Fig. 1D), we concluded that SMPD2 KD does not downregulate LAMP1 proteins through lysosomal degradation.

**Figure 1:**
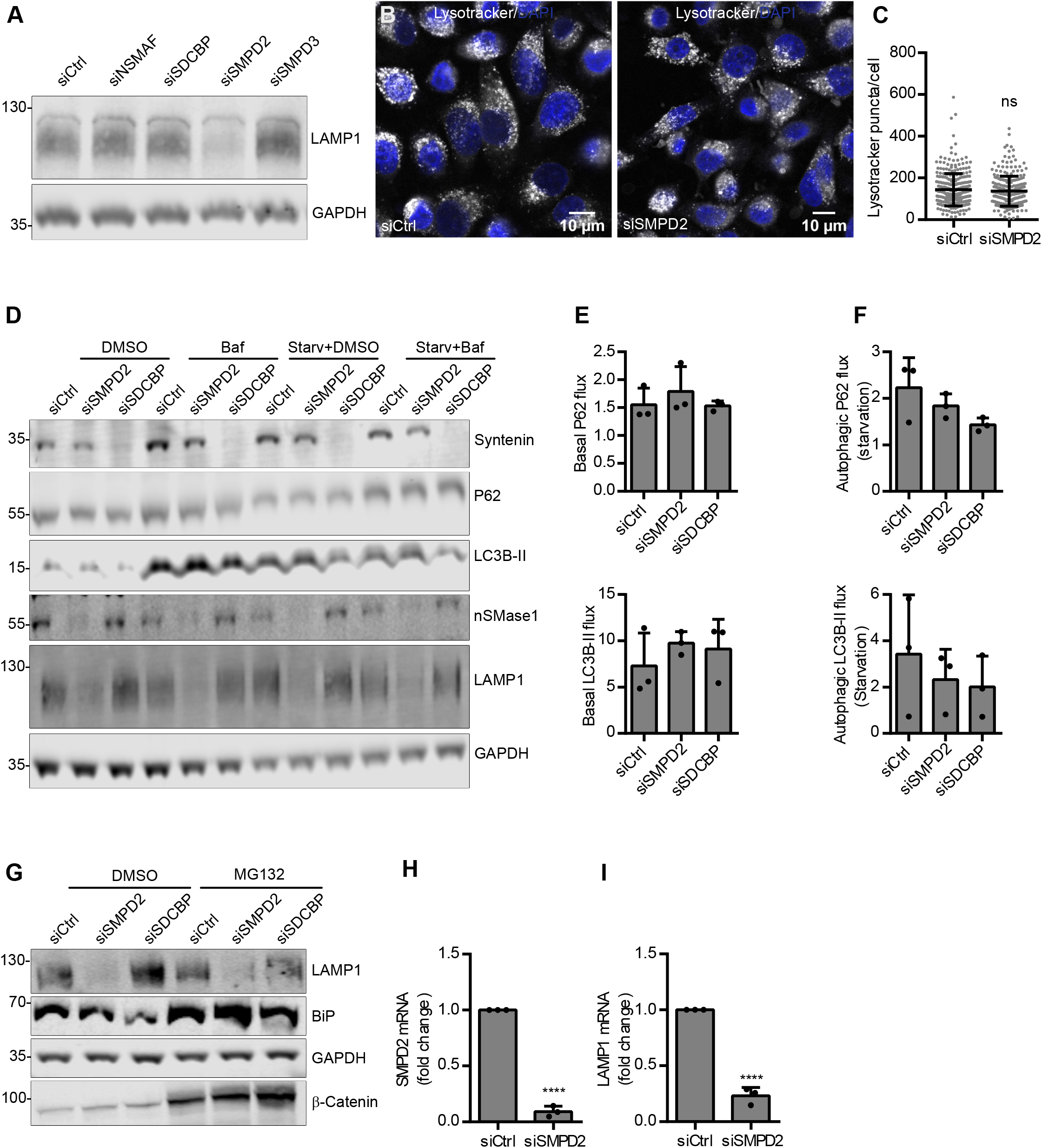
SMPD2 KD downregulates LAMP1 at the mRNA level. **(A)** HeLa cells were transfected with the indicated siRNAs for 48 h and lysates were immunoblotted against LAMP1 and loading control GAPDH. **(B)** HeLa cells transfected with control (Ctrl) and SMPD2 siRNA for 48 h were stained with Lysotracker and DAPI and analyzed by confocal microscopy. **(C)** Quantifications of Lysotracker puncta per cell from (B). Unpaired Student’s two-tailed *t-test*. **(D)** HeLa cells transfected with the indicated siRNAs for 48 h were then grown in FBS containing medium or serum-starved (starv) with or without Bafilomycin A (Baf) for 16 h. **(E)** Quantifications of lysosomal turnover of P62 and LC3B-II under basal conditions from (D). Ordinary one-way ANOVA followed by Dunnett’s multiple comparisons test, not significant. **(F)** Quantifications of lysosomal turnover of P62 and LC3B-II under serum starvation conditions from (D). Ordinary one-way ANOVA followed by Dunnett’s multiple comparisons test. **(G)** HeLa cells transfected with the indicated siRNA for 48 h were then treated with DMSO or MG132 for 16 h. Lysates were immunoblotted against LAMP1, BiP, β-Catenin, and loading control GAPDH. **(H)** Real-time qPCR analysis of SMPD2 mRNA in HeLa cells after 48 h of SMPD2 KD. Unpaired Student’s two-tailed *t-test*, ****p < 0.0001. **(I)** Realtime qPCR analysis of LAMP1 mRNA in HeLa cells after 48 h of SMPD2 KD. Unpaired Student’s two-tailed *t-test*, ****p < 0.0001. All the experiments from (A)-(I) were done in at least three biological replicates.

We next analyzed whether SMPD2 KD downregulates LAMP1 through proteasomal degradation by using proteasomal inhibitor MG132. Efficient proteasomal inhibition was demonstrated by accumulation of β-catenin and ER-resident protein BiP compared with control cells (Fig. 1G), but MG132 did not affect SMPD2 KD-induced LAMP1 downregulation, indicating that SMPD2 KD does not downregulate LAMP1 by proteasomal degradation (Fig. 1G, Fig. S1D). As LAMP1 protein was neither targeted for lysosomal nor proteasomal degradation in SMPD2 KD cells, we next analyzed LAMP1 mRNA levels by qPCR (Fig. 1H). Indeed, the LAMP1 mRNA level was significantly reduced by SMPD2 KD in both HeLa (Fig. 1H) and HCT116 (Fig. S1E) cell lines. In contrast, the mRNA level of LAMP2, another closely related lysosomal membrane protein, remained unchanged upon SMPD2 KD (Fig. S1F). Overall, these data confirm that SMPD2 KD specifically downregulates LAMP1 not via protein degradation pathways but at the mRNA level.

### SMPD2 KD causes inefficient activation of the UPR upon ER stress induction

As a readout for nSMase1 activity, we used a ceramide antibody previously used for sphingomyelin metabolism and ceramide signaling studies (Vielhaber *et al*, 2001; Parashuraman & D’Angelo, 2019; Yabu *et al*, 2015). Confocal microscopy analysis upon SMPD2 KD showed a significant reduction of staining in both mean intensity of the intracellular ceramide puncta and mean intensity of ceramide puncta that colocalize with the ER marker Calnexin (Fig. S1G, H). These data suggest that SMPD2 KD affects cellular ceramide levels specifically at the ER. In addition to the reduced ceramide levels at the ER upon SMPD2 KD (Fig. S1G, H), overexpressed nSMase1 localizes to the ER (Rodrigues-Lima *et al*, 2000; Fensome *et al*, 2000; Tomiuk *et al*, 2000), where transmembrane proteins such as LAMP1 are translated (Schwarz & Blower, 2016). In line with this, we found nSMase1 protein levels significantly increased upon ER stress induction with tunicamycin (Tuni) or thapsigargin (Thapsig) in HeLa cells (Fig. 2A, B). As nSMase1 protein level was increased by ER stress, we next investigated if SMPD2 KD induces ER stress and therefore activates the unfolded protein response (UPR) pathways. ER morphology appeared unchanged upon SMPD2 KD (Fig. S1I). We analyzed three ER stress markers, BiP, Calnexin, and PDI (Protein disulfide-isomerase) which are usually upregulated through UPR pathways upon ER stress induction (Oslowski & Urano, 2011) (Fig. 2C-F). We found BiP to be upregulated at the protein level by both inhibitors, which was slightly impaired by SMPD2 KD after 4 h of treatment (Fig. 2C, D). We next analyzed different markers that are upregulated through UPR activation upon ER stress at the mRNA level. Although SMPD2 KD alone did not activate UPR, SMPD2 KD cells failed to upregulate several ER stress markers (sXBP-1, ATF4, and EDEM) to the same levels as their control counterparts upon stress induction (Fig. 2G-K). Furthermore, under ER stress conditions the LAMP1 mRNA remained significantly downregulated by SMPD2 KD (Fig. 2L). And in alignment with the increased nSMase1 protein levels (Fig. 2A, B), SMPD2 mRNA levels were slightly increased by tunicamycin and thapsigargin treatment (Fig. 2M).

**Figure 2:**
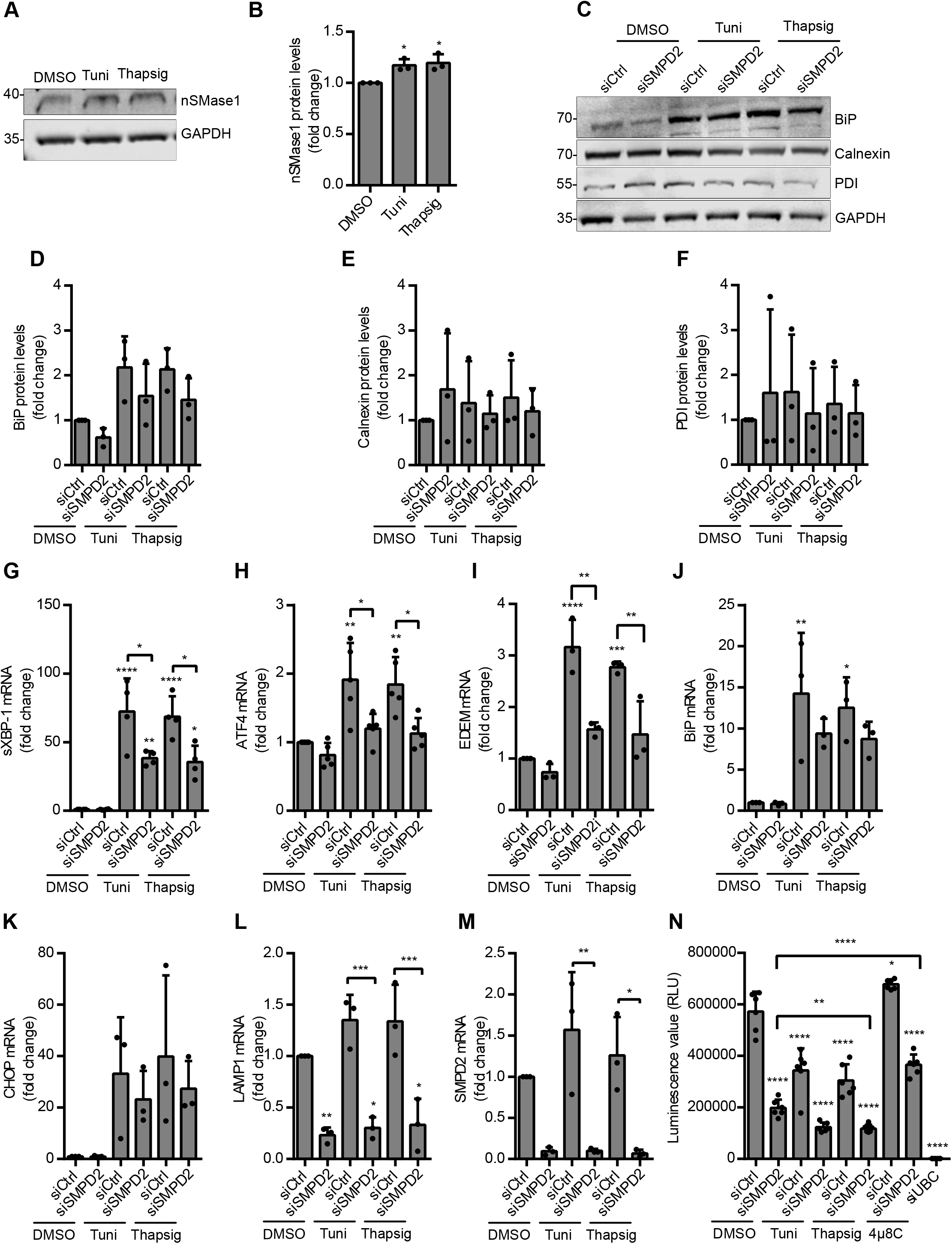
SMPD2 KD causes inefficient activation of ER stress signaling. **(A)** HeLa cells treated with tunicamycin (Tuni) and Thapsigargin for 4 h and cell lysates were immunoblotted against nSMase1 and loading control GAPDH. **(B)** Western blot quantifications of nSMase1 from (A) normalized to the corresponding loading control GAPDH before normalization to the control. Ordinary one-way ANOVA followed by Dunnett’s multiple comparisons test, *p < 0.01 **(C)** HeLa cells transfected with Ctrl or SMP2 siRNA for 48 h were treated with tunicamycin or Thapsigargin for 4 h. Cell lysates were immunoblotted against BiP, Calnexin, PDI, and loading control GAPDH. (D-E) Western blot quantifications of BiP **(D)**, Calnexin **(E)**, and PDI **(F)** normalized to the corresponding loading control GAPDH before normalization to the control. Ordinary oneway Anova followed by Tukey’s multiple comparisons test. (G-M) Real-time qPCR analysis of spliced XBP-1 (sXBP-1) **(G)**, ATF4 **(H)**, EDEM **(I)**, BiP **(J)**, CHOP **(K)**, LAMP1 **(L)**, and SMPD2 **(M)** mRNA in HeLa cells transfected with Ctrl and SMPD2 siRNA for 48 h and treated with tunicamycin or thapsigargin for 4 h. Ordinary one-way Anova followed by Tukey’s multiple comparisons test, ****p < 0.0001, ***p < 0.0009 **p < 0.001, *p < 0.01. **(N)** Viability assay of HeLa cells transfected with Ctrl, SMPD2, and UBC siRNA for 48 h and then treated with 4μ8C (16 h) or tunicamycin or thapsigargin for 4 h. Ordinary oneway Anova followed by Tukey’s multiple comparisons test, ****p < 0.0001, ***p < 0.0009 **p < 0.001, *p < 0.01. siRNA against UBC was used as a negative control for the viability assay. All experiments above, unless otherwise stated, are done in three biological replicates.

The UPR activation upon ER stress results in mRNA degradation through IRE-1 RNase activity (the RIDD pathway) to relieve stress by reducing protein translation burden in the ER (Hollien *et al*, 2009). LAMP1 mRNA was identified as a substrate of IRE-1-dependent decay of mRNAs (Maurel *et al*, 2014). If LAMP1 mRNA is degraded through the RIDD pathway upon SMPD2 KD, we hypothesized that IRE-1 RNase activity inhibitor 4μ8C (Cross *et al*, 2012) should restore LAMP1 mRNA and protein levels. IRE-1 cleaves XBP-1 (X-box binding protein) mRNA into two transcripts, known as spliced XBP1 and 2 (sXPB1 and 2) (Maurel *et al*, 2014). The efficient inhibition of the IRE-1 RNase activity by 4μ8C was confirmed by the absence of spliced XBP-1 (sXBP-1) mRNA (Fig. S1J). However, LAMP1 mRNA, as well as protein levels, remained downregulated, when cells were treated with 4μ8C overnight after 48 h of SMPD2 KD (Fig. S1K, L). Surprisingly, even under ER stress conditions induced by Tunicamycin treatment (Fig. S1L) SMPD2 KD-downregulated LAMP1 protein level did not recover under 4μ8C treatment. These data indicate that reduced mRNA level upon SMPD2 KD is not caused by degradation through the RIDD pathway.

Since SMPD2 KD cells failed to activate an efficient UPR signaling upon ER stress induction and are less viable under normal conditions, we next analyzed how SMPD2 KD affects cellular fitness under ER stress. Indeed, SMPD2 KD cells are less viable upon ER stress than their control counterparts; especially, SMPD2 KD cells treated with thapsigargin are significantly less viable than SMPD2 KD treated with DMSO. Surprisingly, 4μ8C significantly increased viability in both control and SMPD2 KD cells (Fig. 2N). In summary, the above data suggest that although SMPD2 KD does not downregulate LAMP1 mRNA through the RIDD pathway and does not induce ER stress, a full-potential UPR signaling activation upon ER stress is not achieved in SMPD2 KD cells, and consequently their cellular fitness is impaired under ER stress conditions.

### SMPD2 KD arrests cells in the G1 phase

As SMPD2 KD significantly decreased cell viability (Fig. S1A, 2N), and JNK signaling was shown to induce apoptosis via nSMase1-induced ceramide generation under various stress conditions (Yabu *et al*, 2015), we next investigated whether SMPD2 KD induces apoptosis using flow cytometry (Fig. S2A). In alignment with the previous results (Fig. S1A, 2N), SMPD2 KD significantly reduced viable cells in an Annexin/PI assay (Fig. 3A). Although early apoptosis was significantly increased (A^-^/PI^+^), no significant change in late apoptosis (A^+^/PI^+^) and necrosis levels (A^-^/PI^+^) were observed upon SMPD2 KD (Fig. 3A). Concordantly, SMPD2 KD did not induce the cleavage of different apoptosis marker proteins as determined by western blot analysis of Caspase-3, −7, and PARP (Fig. 3B, Fig.S2B-E).

**Figure 3:**
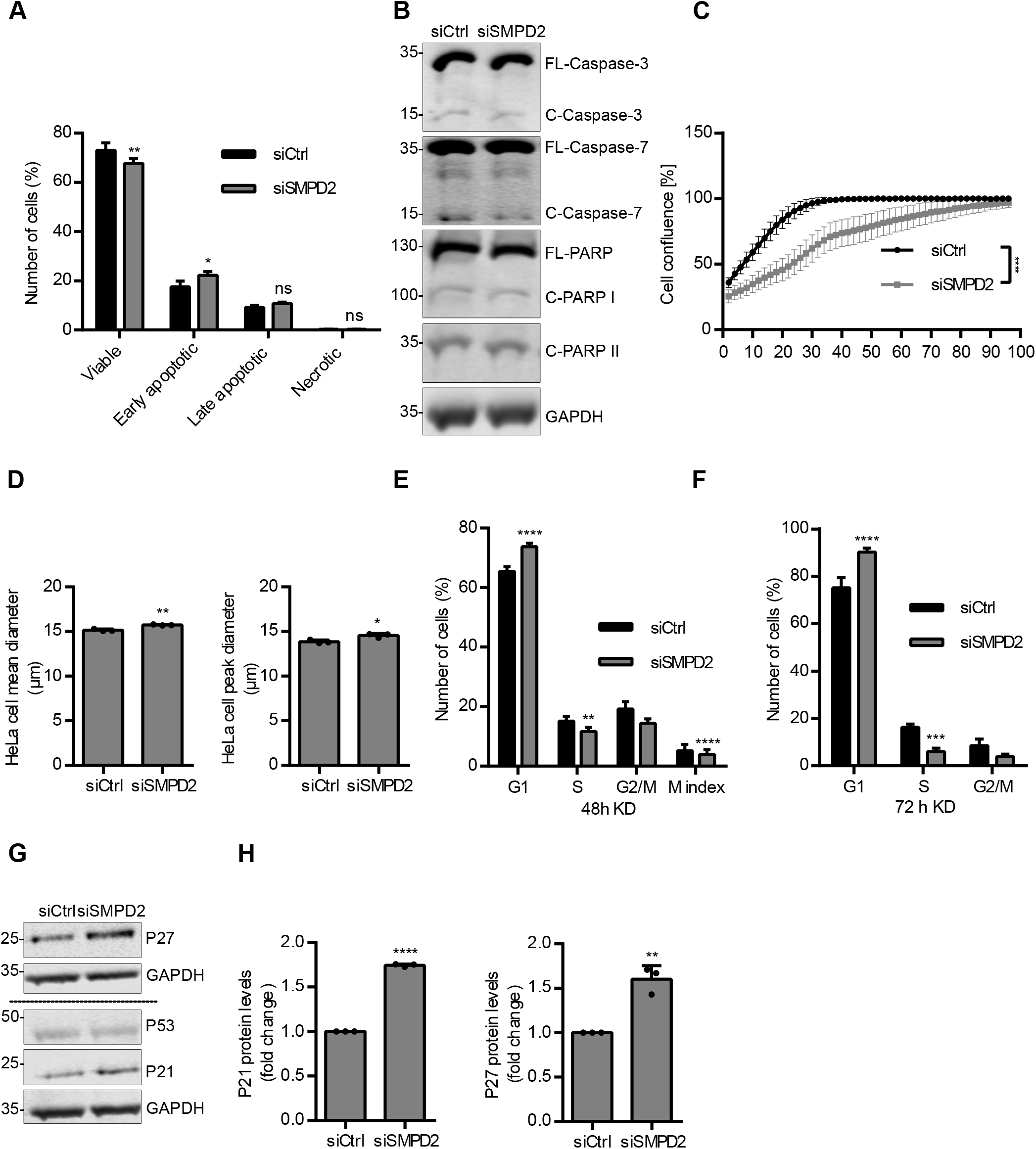
SMPD2 KD arrests cells in the G1 phase. **(A)** Percentage of HeLa cells transfected with Ctrl or SMPD2 siRNA for 48 h were analyzed by flow cytometry and sorted into Viable (Annexin-V^-^/PI^-^), Early apoptotic (Annexin-V^+^/PI^-^), Late apoptotic (Annexin-V^+^/PI^+^), and Necrotic (Annexin-V^-^/PI^+^). Twoway ANOVA followed by Sidak’s multiple comparisons test, **p < 0.001, *p < 0.01. **(B)** HeLa cells were transfected with Ctrl or SMPD2 siRNA for 48 h and then cell lysates were immunoblotted against different apoptosis markers and loading control GAPDH. **(C)** HeLa cells transfected with Ctrl or SMPD siRNA for 48 h were replated and seeded into a 96-well plate for Incucyte proliferation analysis. Proliferation rate was measured by cell confluence percent versus time by taking images of the wells at 2 h intervals for the indicated duration. Paired Student’s two-tailed *t-test*, ****p < 0.0001. **(D)** HeLa cell mean diameter (Left) and peak diameter (Right) determined with Casy cell counter after 72 h of Ctrl or SMPD2 siRNA transfection. Unpaired Student’s two-tailed *t-test*, *p < 0.01. (E-F) Percentage of HeLa cells transfected with Ctrl or SMPD2 siRNA for 48 h **(E**) and 72 h **(F)** were gated for G1, S, and G2/M. Mitotic index in (E) was determined from the G2/M population positive for p-Mpm-2 staining. Two-way ANOVA followed by Sidak’s multiple comparisons test, ****p < 0.0001, ***p < 0.0009 **p < 0.001. **(G)** Lysates from HeLa cells transfected with Ctrl or SMPD2 siRNA for 72 h were immunoblotted against P27, P53, P21, and loading control GAPDH. **(H)** Western blot quantifications of P21 (Left) and P27 (Right) normalized to the corresponding loading control GAPDH before normalization to the control. Unpaired Student’s two-tailed *t-test*, ****p < 0.0001, **p < 0.001. All experiments from (A) to (H) are done in three biological replicates.

Since the reduced cell viability upon SMPD2 KD is not caused by apoptosis, we next analyzed the cell proliferation rate by incucyte imaging. Indeed, the cell proliferation rate was significantly lowered by SMPD2 KD (Fig. 3C). Additionally, SMPD2 KD cells are significantly bigger when compared to the control cells (Fig. 3D, Fig. S2F), suggesting an SMPD2 KD-induced cell cycle delay. Cell cycle progression analysis by flow cytometry (Fig. S2G) revealed SMPD2 KD to significantly increase the percentage of cells in G1 (Fig. 3E). The percentage of cells in G1 was furthermore increased (by around 20 %) when SMPD2 KD duration was increased from 48 h to 72 h (Fig. 3F) and reduced slightly after 96 h when compared to the 72 h SMPD2 KD (Fig. S2H). Overall, these data indicate that a significant number of SMPD2 KD cells show a delay in cell cycle progression. To test whether SMPD2 KD causes a transient G1 arrest, we analyzed the protein expression of two G1 cell cycle arrest markers – P21 and P27, along with P53, which regulate these two proteins (Chen, 2016). SMPD2 KD significantly upregulated the protein expression of both P21 and P27 (Fig. 3G, H), while P53 protein levels remained unchanged upon SMPD2 KD (Fig. S3A). In agreement with the FACS data (Fig. 3F), both P21 and P27 protein expression reached a maximum after 72 h of SMPD2 KD (Fig. 3G, H) when compared to 96 h SMPD2 KD (Fig. S3B, C). In summary, these data indicate that SMPD2 KD cells are bigger and proliferate slower than the control cells, as they are temporarily arrested in the G1 phase of the cell cycle at the peak of the transient SMPD2 KD.

### SMPD2 KD downregulates the Wnt signaling pathway

Next, we investigated molecular pathways through which SMPD2 KD causes G1 cell cycle arrest. Since various lipids in the nucleus are indicated to regulate many processes including gene transcription (Martelli *et al*, 2004), we analyzed if SMPD2 KD upregulates P21 and P27 for G1 arrest through the P53-dependent DNA damage response pathway. DNA damage, through ataxia telangiectasia mutated (ATM) or Rad3-related protein (ATR), phosphorylates two checkpoint kinases (Chk1 and Chk2), which then activates P53 by phosphorylation (Chen, 2016) (Fig. 4A). Activated P53 upregulates, among many of its target genes, two cell cycle inhibitory proteins P21 and P27. Therefore, an increase in phosphorylated Chk1 and Chk2 could indicate activation of the DNA damage response pathway upon SMPD2 KD. The expression level of the phosphorylated Chk1 and phosphorylated Rb (Retinoblastoma protein), another cell cycle inhibitory protein, was not changed by SMPD2 KD (Fig. 4B, C; Fig. S3D). However, SMPD2 KD significantly decreased the phosphorylated Chk2 level (Fig. 4B, C), in line with a role of Chk2 at the crossroad of DNA damage and metabolic fitness (Ajazi *et al*, 2021).

**Figure 4:**
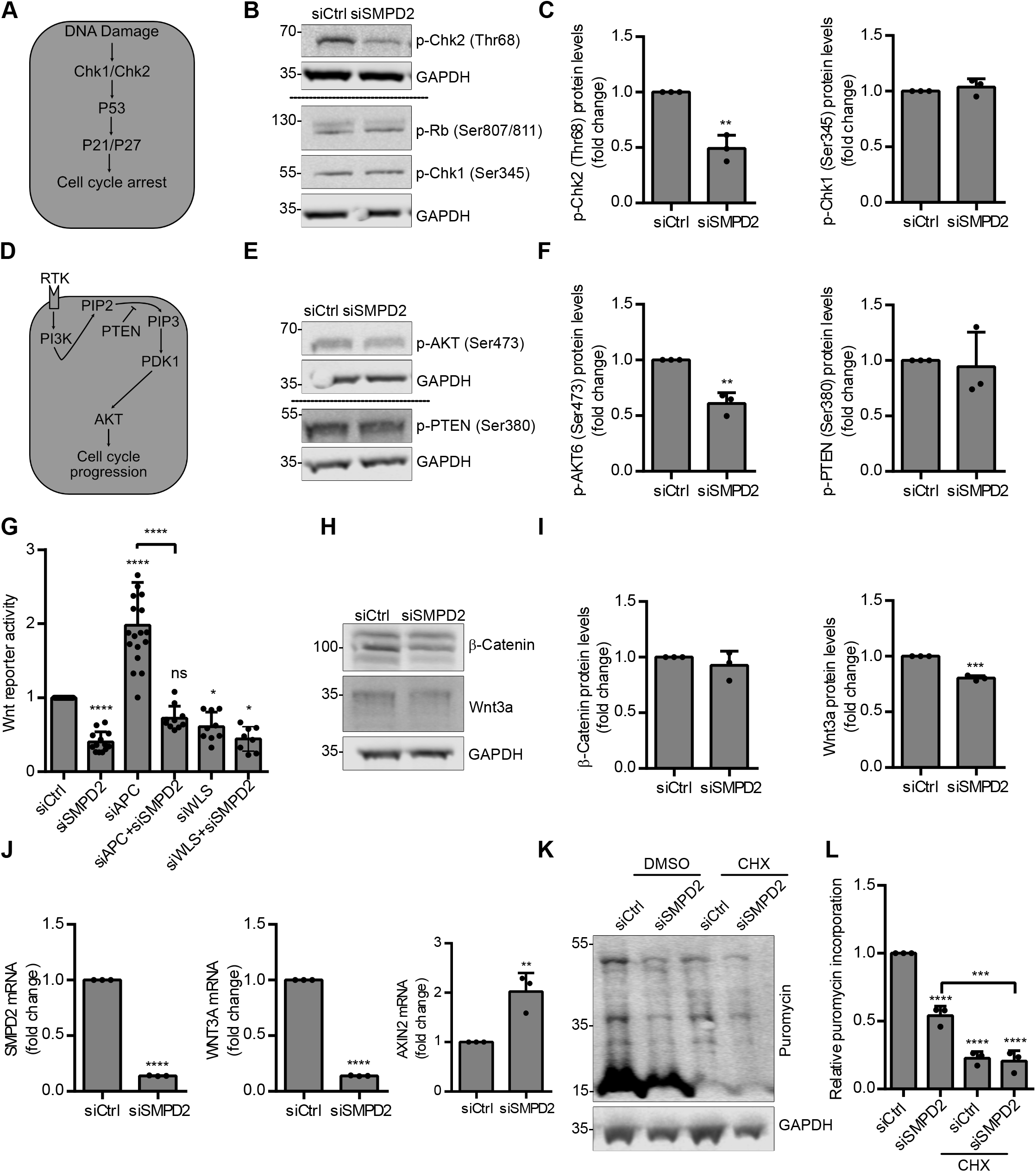
Wnt signaling is downregulated in SMPD2 KD cells. **(A)** Scheme showing P53-dependent DNA damage response pathway that mediates G1 cell cycle arrest. **(B)** Lysates from HeLa cells transfected with Ctrl or SMPD2 siRNA for 48 h were immunoblotted against the p-Chk2, p-Chk1, p-Rb, and loading control GAPDH. **(C)** Western blot quantifications of p-Chk1 and p-Ch2 from (B) normalized to the corresponding loading control GAPDH before normalization to the control. Unpaired Student’s two-tailed *t-test*, **p < 0.001. **(D)** Scheme showing PI3K/Akt signaling pathway. (**E)** Lysates from HeLa cells transfected with Ctrl or SMPD2 siRNA for 48 h were immunoblotted against the p-Akt, p-PTEN, and loading control GAPDH. **(F)** Western blot quantifications of p-Akt and p-PTEN from (E) normalized to the corresponding loading control GAPDH before normalization to the control. Unpaired Student’s two-tailed *t-test*, **p < 0.001. **(G)** Wnt reporter activity analysis in HCT116 cells transfected with the indicated siRNAs for 72 h measured using TCF/LEF luciferase reporter assay. The reporter vectors were transected 24 h after siRNA reverse transfection. Ordinary one-way Anova followed by Tukey’s multiple comparisons test, ****p < 0.0001, *p< 0.01. **(H)** HCT116 cells were transfected with Ctrl or SMPD2 siRNAs for 72 h and lysates were immunoblotted against β-catenin, Wnt3a, and loading control GAPDH. **(I)** Western blot quantifications from (I) normalized to the corresponding loading control GAPDH before normalization to the control. Unpaired Student’s two-tailed *t-test*, ***p < 0.0009. (H-L) **(J)** Real-time qPCR analysis of SMPD2, WNT3A, and AXIN2 mRNA in HCT116 cells transfected with Ctrl and SMPD2 siRNA for 72 h. Unpaired Student’s two-tailed *t-test*, ****p < 0.0001. **(K)** HeLa cells were transfected with Ctrl or SMPD2 siRNA for 48 h and treated with or without Cycloheximide (CHX) for 16 h. Lysates, after labeling with puromycin for 1 h, were immunoblotted against puromycinylated proteins with puromycin antibody. **(L)** Western blot quantifications from (K) normalized to the corresponding loading control GAPDH before normalization to the control. Ordinary one-way Anova followed by Tukey’s multiple comparisons test, ****p < 0.0001, ***p < 0.009. All experiments from (A)-(L) are done in three biological replicates.

Extracellular growth signals are integrated into intracellular signals by pathways such as PI3K/Akt to drive cell cycle progression (Vivanco & Sawyers, 2002) (Fig. 4D). SMPD2 was shown to exhibit a negative gene interaction with PTEN (phosphatase and tensin Homolog), a tumor suppressor gene that inhibits G1-S transition by antagonizing PI3k/Akt pathway activation (Fig. 4D) (Vizeacoumar *et al*, 2013). Therefore, we analyzed the level of phosphorylated PTEN and AKT upon SMPD2 KD. Although phosphorylated PTEN levels remained unchanged, SMPD2 KD significantly reduced the level of phosphorylated Akt (Fig. 4E, F) in line with a role of SMPD2 KD in G1-S transition through the PI3K/Akt pathway.

Wnt signaling is another major pathway that drives cell proliferation (Freese *et al*, 2010). Interestingly, SMPD2 KD reduced basal Wnt activity, but also Wnt activity induced by β-catenin accumulation through APC KD (Fig. 4G). Reduction of APC KD-induced Wnt activity by SMPD2 KD indicates that the KD affects the Wnt signaling at the level or downstream of β-catenin. Indeed, β-catenin protein levels were downregulated by SMPD2 KD in HeLa cells (Fig. S3E, F). Surprisingly, in HCT116 cells, although β-catenin protein levels remained unchanged, Wnt3a protein levels were significantly reduced by SMPD2 KD (Fig. 4H, I). We furthermore analyzed Wnt signaling components upon SMPD2 KD by qPCR in HCT116. In alignment with the reduction in Wnt3a protein levels, SMPD2 KD dramatically reduced Wnt3a mRNA (Fig. 4J) and significantly upregulated AXIN2 mRNA (Fig. 4J), which is involved in the negative feedback loop of Wnt signaling (Lustig *et al*, 2001). The mRNA levels of the other Wnt signaling components CTNNB-1, EVI, and WNT7B remained unchanged upon SMPD2 KD (Fig. S3G). Additionally, SMPD4, another member of nSMases, showed no compensatory increase in mRNA levels upon SMPD2 KD (Fig. S3H). Since many proteins and mRNA are downregulated upon SMPD2 KD, we next analyzed the overall protein translation upon SMPD2 KD by puromycin assay (Goodman & Hornberger, 2013) and using protein translation inhibitor cycloheximide (CHX) as a negative control. Indeed, SMPD2 KD significantly reduced the overall protein translation (Fig. 4K, L).

In summary, the data above indicate that SMPD2 KD seems to affect many cellular processes by downregulating their signaling components including Wnt signaling – which could explain the reduction in global protein translation and G1 cell cycle arrest.

## Discussion

We found that nSMase1 plays an important role in maintaining cellular fitness. NSMase1 is required for activating a full-potential adaptive UPR upon ER stress induction with tunicamycin and thapsigargin-two inhibitors that cause ER stress through distinct mechanisms. SMPD2 KD cells failed to upregulate multiple genes that are usually upregulated through UPR pathways upon ER stress to the same levels as their control counterparts. Furthermore, we show that nSMase1 is necessary for proper cell cycle progression during the G1 phase as SMPD2 KD led to a significant G1 arrest. Moreover, the overall protein translation at the ribosome was reduced by SMPD2 KD. These data, for the first time, establish an important biological role for SMPD2 at the level of UPR activation upon ER stress and cell cycle progression.

We further show that nSMase1 regulates LAMP1 protein expression at the level of its mRNA but not LAMP2. LAMP1 and LAMP2 are the major glycoproteins localized on the lysosomal membrane and thought to contribute to lysosomal integrity, catabolism, and pH maintenance (Sawada *et al*, 1993). SMPD2 KD downregulated LAMP1 mRNA and thereby reduced its protein expression. Interestingly, lysosomal acidification and function remain unaffected by this significant reduction in the LAMP1 protein levels as both lysotracker staining and the autophagy flux remained unchanged. Therefore, the downstream biological implication of this dramatic LAMP1 downregulation upon SMPD2 KD would be of great interest for future studies.

ER stress is sensed by three ER membrane proteins – IRE1α, PERK, and ATF6α – which initiate UPR through three distinct pathways by upregulating different proteins (Hetz *et al*, 2020). Although nSMase1 expression was increased by ER stress, SMPD2 KD alone did not induce an ER stress condition as no UPR activation was observed. Upon ER stress induction, SMPD2 KD cells failed to upregulate multiple genes under the control of all three master regulators. These data indicate that SMPD2 KD affects all the three UPR pathways in the ER. Indeed, several early studies showed that overexpressed nSMase1 resides in the ER (Fensome *et al*, 2000; Rodrigues-Lima *et al*, 2000; Tomiuk *et al*, 2000) and recent affinity-based mass spectrometry studies showed that nSMase1 interacts with multiple ER-resident proteins including reticulon 3 and reticulon 4 (Huttlin *et al*, 2017, 2021). Therefore, based on the observations from our study, it is clear that nSMase1 plays a functional role in restoring ER function and homeostasis under ER stress conditions. The UPR signaling activation persists until an adaptive response to the ER stress is achieved, in the case of unresolvable stress, the UPR turns pro-apoptotic to induce cell death (Hetz *et al*, 2020). Indeed, due to their inability to mount a full-potential UPR signaling, SMPD2 KD cells are less viable upon ER stress induction.

Additionally, SMPD2 KD significantly downregulated LAMP1 and Wnt3a mRNAs – both of which are translated and subsequently modified in the ER as an integral membrane and a secretory protein respectively. NSMase1 could affect signaling recognition particle (SRP)-dependent targeting of these proteins to the ER for translation as nSMase1 was shown to interact with both signal recognition particle receptor B (SRPRB) and the translocon-associated signal sequence receptor 2 (SSR2). These two proteins are ER membrane proteins required for proper docking of the whole mRNA:ribosome:polypeptide:SRP complex to the ER for efficient translation (Akopian *et al*, 2013). It is therefore conceivable that nSMase1 plays an important role in the translation of these proteins possibly by providing specific lipids to establish membrane nanodomains for efficient docking of this co-translation machinery to the ER. In this way, downregulation of LAMP1 and Wnt3a could be due to a disrupted SRP-dependent targeting of these mRNAs to the ER for translation upon SMPD2 KD. In line with its ER localization and potential functions, nSMase1 also interacts with several GPI (Glycosylphosphatidylinositol) anchor attaching proteins such as glycosylphosphatidylinositol anchor attachment 1 (GPAA1) and Phosphatidylinositol glycan anchor biosynthesis, class S (PIG-S) (Huttlin *et al*, 2021).

Overall, the data from others and our study demonstrate that nSMase1 at the ER plays an important role in activating an efficient adaptive UPR upon ER stress and could also regulate other aspects of ER function such as translation and post-translational modifications.

Since SMPD2 KD cells are arrested in the G1 phase with both P21 and P27 upregulated, it is possible that nSMase1 activity, by generating required structural lipids, might drive the remodeling of the nuclear envelope, chromatin, and nuclear matrix during cell cycle progression (Mizutani *et al*, 2001). Alternatively, since nSMase1 belongs to the superfamily of endo/exonucleases (Bill X.Wu, Christopher J.Clarke, 2010) that cleave the phosphodiester bond between the successive nucleotides, it may hydrolyze the phosphodiesterase bond of the DNA. As described for many endo/exonucleases (Nishino & Morikawa, 2002), it, therefore, could be involved in DNA repair and cell cycle progression. In this way, inefficient DNA repair upon SMPD2 KD might induce a DNA-damage-like response that upregulates P21 and P27 to induce G1 cell cycle arrest. Therefore, it is noteworthy to further investigate the activation of tumor suppressor gene P53 upon SMPD2 KD. P53 senses DNA damage and mediates cell cycle arrest by upregulating P21 and P27 (Chen, 2016).

Duplication of not only cellular DNA content but also of membranes and organelles is a prerequisite for cellular growth and division. Lipids – both storage and membrane – play an important role in cell cycle regulation (Storck *et al*, 2018). For example, two triacylglycerol lipases Tgl3 and Tgl4 are required for efficient cell cycle progression during the G1/S transition in yeast cells (Chauhan *et al*, 2015). Specifically, lipolysis-derived sphingolipids activate PP2A, a major cell cycle regulating protein phosphatase, to dephosphorylate SWE1 (human ortholog WEE1) for efficient cell cycle progression (Chauhan *et al*, 2015). Therefore, it is conceivable that ceramide generated by nSMase1 acts as a second messenger targeting proteins involved in cell-cycle regulation such as PP2A. Indeed, several studies proposed PP1 and PP2A as the potential intracellular protein target of ceramide. PP2A dephosphorylates over 300 substrates involved in the cell cycle, thereby regulating almost all the major pathways including the Wnt pathway and the cell cycle checkpoints (Wlodarchak & Xing, 2016). Therefore, reduced Wnt-signaling could be also due to deregulated PP2A activity upon SMPD2 KD. Collectively, these data suggest that nSMase1 plays an important role in overall cellular fitness and proliferation without which cells enter a transient G1 arrest with reduced global protein translation.

So far, the biological role as well the physiological substrate for nSMase1 is unknown. Our study shows that nSMase1 plays a vital role in overall cellular fitness and survival as SMPD2 KD induced transient G1 arrest. Additionally, nSMase1 activity is essential for activating an efficient UPR upon ER stress induction. The observations and data described herein would serve as a strong foothold for further studies in unraveling not only the molecular function but also the molecular pathways through which nSMase1 contributes to maintaining cellular homeostasis.

## Materials and methods

### Cell culture and transfection (siRNA and plasmids)

HeLa (kindly provided by Holger Bastians, Goettingen) and HCT116 cells (DSMZ) were maintained in DMEM (Thermo Scientific Life technologies) supplemented with 10% fetal calf serum (Biochrom) and 10 μg/ml Penicillin/Streptomycin (Sigma-Aldrich) at 37 °C in a humidified atmosphere with 5% CO_2_. Cells were transiently transfected with Screenfect siRNA (Dharmacon) for indicated duration according to the manufacturer’s instructions. The cells were authenticated and routinely checked for mycoplasma contamination. Cells were treated with the drugs in the following concentrations for indicated durations. Bafilomycin (Sigma-Aldrich, 100ng/ml, 16 h); MG132 (Sigma-Aldrich, 100ng/ml, 16 h); 4μ8C (Millipore Merck, 49 μM, Duration as indicated); Cycloheximide (Carlroth, 20 μg/ml, 16 h).

**Table 1:**
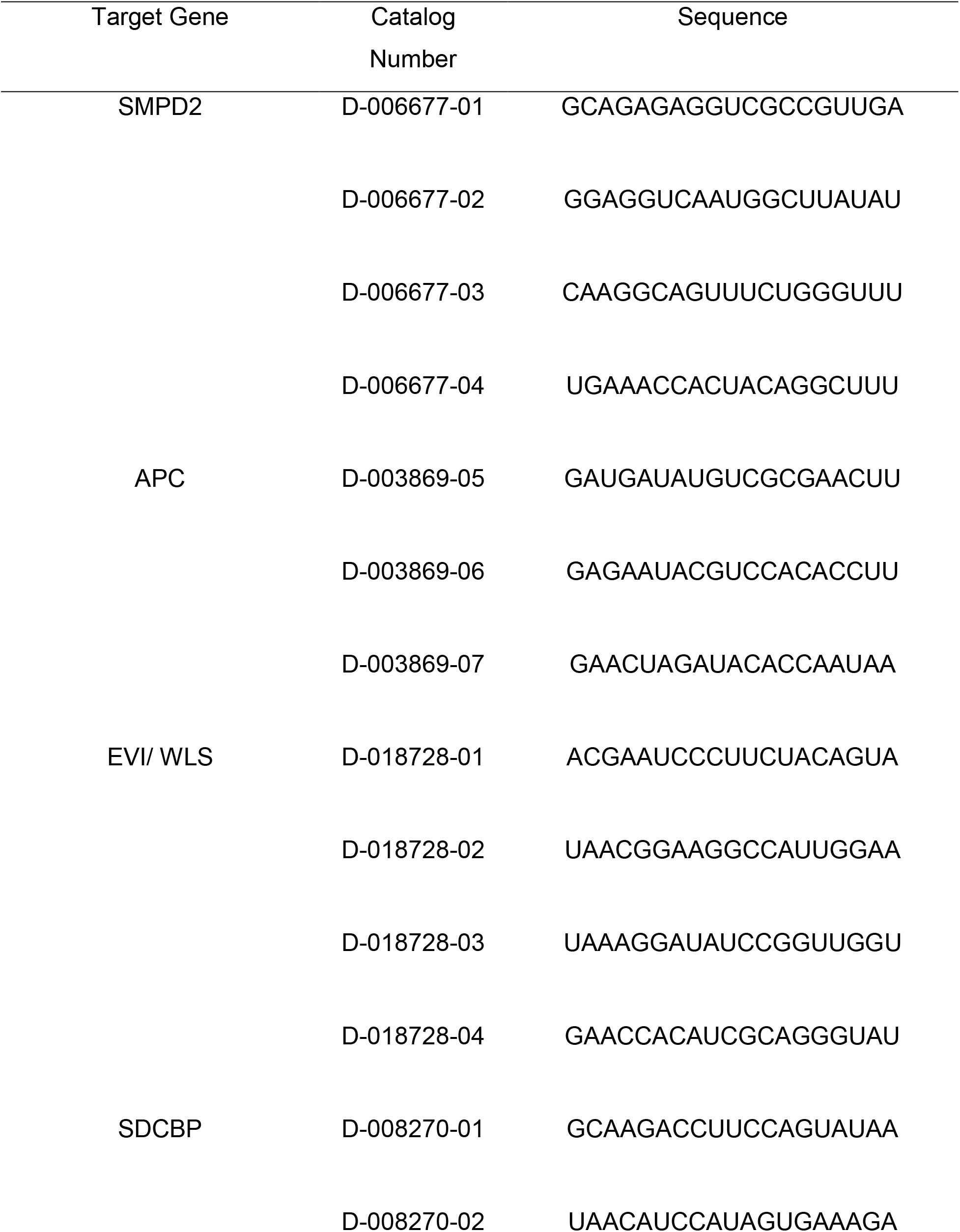
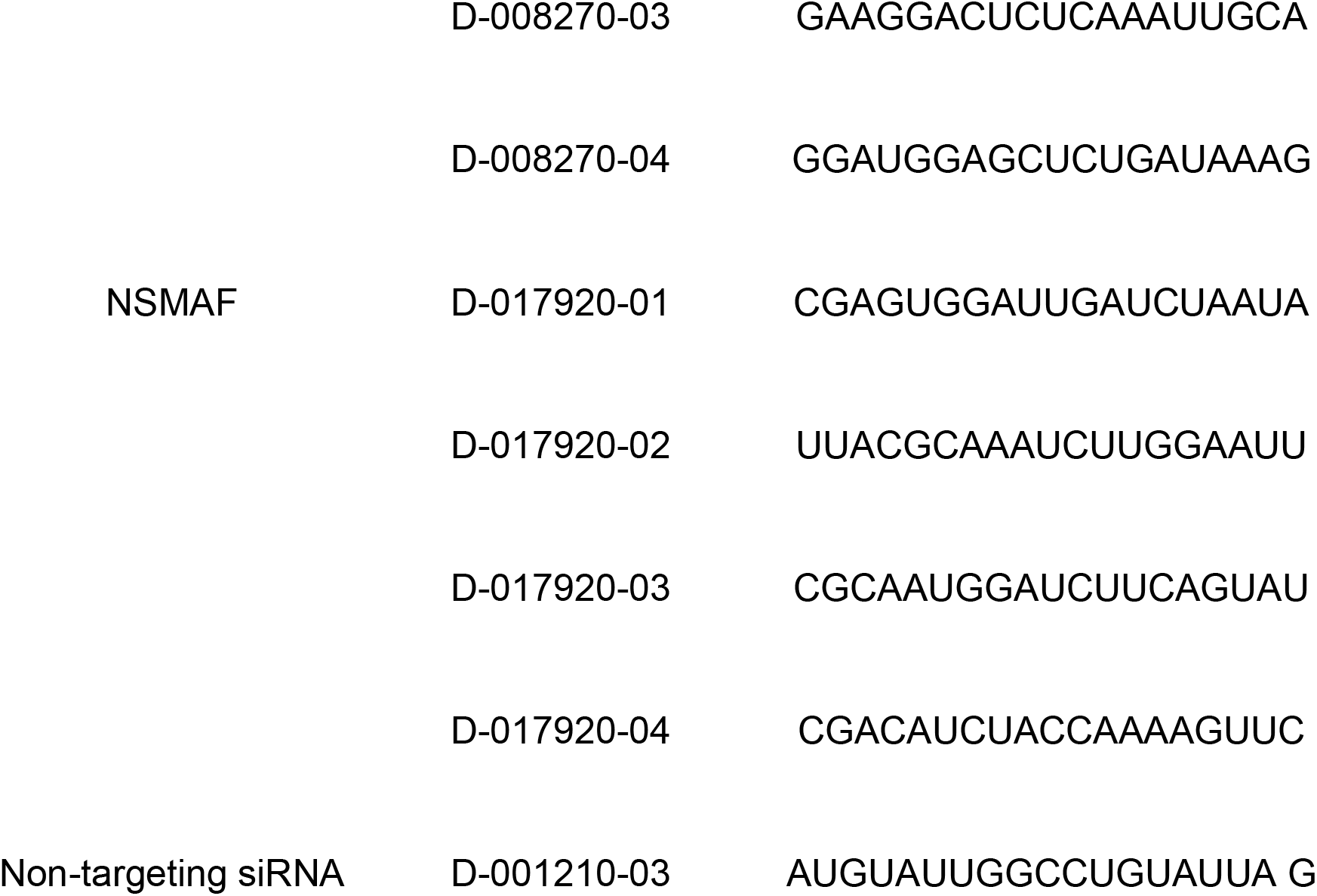
Dharmacon siRNA SMARTpools against the indicated genes used in the study

### Antibodies

The following antibodies and dilutions were used for western blot (WB) or for immunofluorescence staining (IF): LAMP1 (WB 1:1000, Abcam, ab24170); Syntenin (WB 1:1000, Abcam, ab133267); GAPDH (WB 1:1000, Millipore, CB1001); Calnexin (WB 1:1000, Abcam, ab75801), P62 (WB 1:1000, Abnova); LC3B (WB 1:1000, Santa Cruz); Ceramide (IF 1:100, Enzo, ALX-804-196); BiP (WB 1:1000, CST), nSMase1 (IF 1:100, WB 1:1000, Sigma); β-Catenin (WB 1:1000, BD Transduction), PDI (WB 1:1000, CST), FL-Caspase 3 (WB 1:1000, CST); C-Caspase-3 (WB 1:100, CST), FL-Caspase 7 (WB 1:1000, CST); C-Caspase-7 (WB 1:1000); FL-PARP (WB 1:1000, CST), C-PARP (WB 1:1000, CST), p-CHk2 (Thr68) (WB 1:1000, CST); p-Rb (Ser807/811) (WB 1:1000, CST); p-Chk1 (Ser345) (WB 1:1000, CST); p-AKT (Ser473) (WB 1:1000, CST); Puromycin (WB 1:1000, Millipore);; pPTEN (Ser380) (WB 1:1000, CST)

### Western blot analysis

Cell lysates in the SDS-PAGE sample buffer were boiled for 5 min before separating the protein on 4-12% gradient gels (Bolt Bis-Tris Plus Gels, Thermo Scientific). Proteins were then transferred onto PVDF membranes (Merck) and blocked with 5% (wt/vol) milk-TBST for 30 min before incubating with primary antibodies overnight at 4°C. After washing, membranes were incubated with fluorescently labeled secondary antibodies at room temperature in the dark and detected using the Li-COR Odyssey system. Western blot quantifications were done with Odyssey Infrared Image Studio Lite Ver 5.2

### Autophagy flux measurement

Densitometric analysis of the LC3B-II, P62, and GAPDH bands was performed using Odyssey Infrared Image Studio Lite Ver 5.2. To determine autophagy flux, under both basal and serum-starvation autophagy induced conditions, lysosomal turnover of endogenous LC3B-II and P62 were determined. LC3B-II and P62 bands were normalized to GAPDH (LC3B-II/GAPDH, P62/GAPDH). Autophagy flux was then determined by dividing the normalized value of the Bafilomycin-treated lysate by the normalized value of the control-treated lysate of the same sample(Tanida *et al*, 2005).

### Immunostaining, microscopy, and image analysis

Cells were grown in 8-well microscopic coverslips (Sarstedt and IBIDI for live imaging), reverse transfected with indicated siRNAs, and were fixed and permeabilized with 4% paraformaldehyde and 0.2 % Triton-X-100. The slides were then blocked in 3 % bovine serum albumin diluted in PBS, followed by 90 min incubation with primary antibodies. After washing three times, cells were incubated with secondary antibodies conjugated to Alexa-Fluor-488 or 546. Nuclei and actin were labeled by Hoechst and conjugated phalloidin respectively. 200 nM ER tracker (Thermo Scientific Life technologies, E34250) were added to live cells. The cells were visualized with a Zeiss LSM780 confocal microscope (Plan Neofluor 63X/oil NA 1.4 objective). Staining and microscopy conditions were kept identical for comparisons. The number of puncta/cell quantifications were performed using available pipelines with some modifications in CellProfiler (Broad Institute of MIT and Harvard).

### CellTiter-Glo^®^ luminescent viability assay

Cell viability was measured by performing a CellTiter-Glo assay (Promega, G8461). Cells were seeded in a 96-well plate. After reverse siRNA transfection or drug treatment for the indicated duration, 100 *μ*l of the cell titer glow reagent (Promega) diluted to 1:5 with PBS were added to each well. The plate was incubated on a shaker for 2 min at RT to allow cell lysis and then incubated without shaking for 10 min at RT to allow luminescence signal stabilization. The signal was measured using a Centro LB 960 microplate luminometer (Berthold Technologies) and data were analyzed using MikroWin 2000 lite Version 4.43.

### Incucyte proliferation assay

HeLa cells transfected with control or SMPD2 siRNA for 48 h were replated and 6000 transfected cells per well were seeded into Incucyte^®^ Imagelock 96-well plates. The realtime proliferation of the cells was detected by the Incucyte® S2 Live Cell Analysis System (Sartorious) by imaging each well at 2 h intervals.

### RNA isolation and real-time qPCR

Total cellular RNA was isolated using Trizol (Thermo Scientific Invitrogen). The reverse transcription was carried out with 2 μg of RNA. The resulting cDNA product was analyzed by real-time quantitative PCR using Taq Universal SYBRgreen Supermix (Bio-Rad). Transcript Ct-values were converted to fold change expression changes (2-ΔΔCt values) after normalization to the housekeeping gene β-actin. Quantitative real-time PCR was performed using a CFX system (Bio-Rad).

**Table 2:**
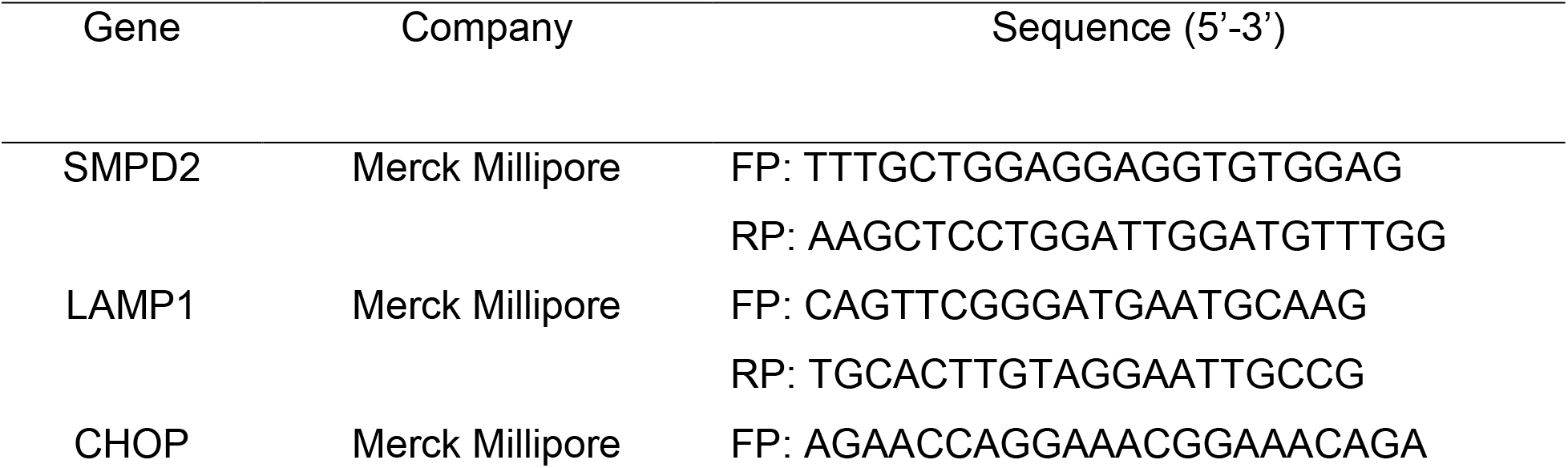
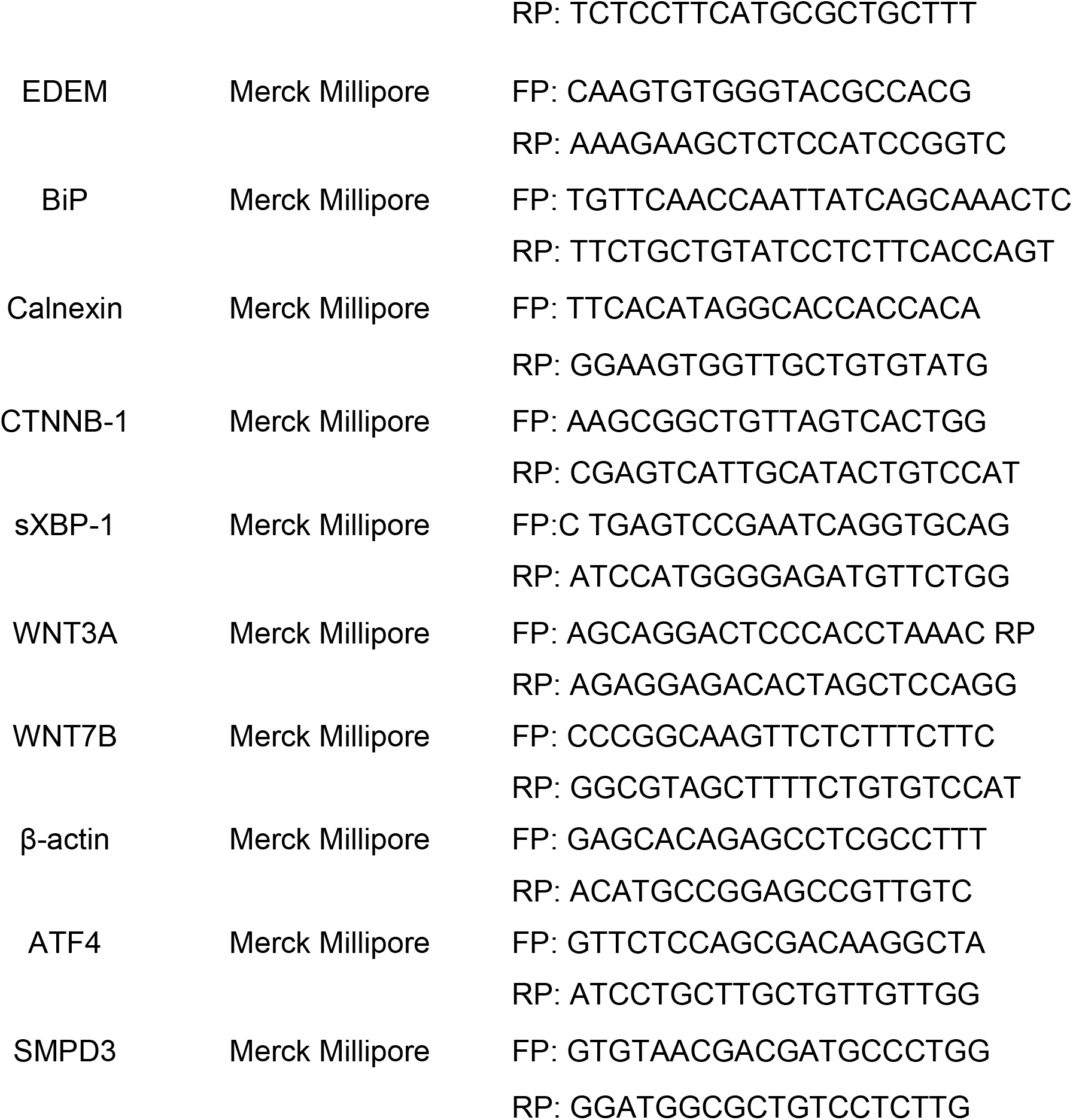
List of primers with their respective sequences used in the study

### Wnt reporter activity assay

Endogenous Wnt signaling activity was assessed by employing a dual-luciferase β-catenin/TCF4 reporter assay. HCT116 cells were transfected with a TCF firefly reporter plasmid (pgGL4.1-6xKD FLuc) (Demir *et al*, 2013) and a constitutively β-catenin renilla reporter plasmid (pgGL4.1-RLuc) (Demir *et al*, 2013) 24 h after reverse transfection with the indicated siRNA. Relative Wnt reporter activity was determined by normalizing the firefly luminescence value to the renilla luminescence value after 48 h of the plasmid transfection.

### Cell cycle analysis by flow cytometry

Flow cytometry was used to analyze cell cycle distribution. Trypsinized cells pelleted at 400 g for 5 min were fixed by adding cold 70% ethanol dropwise while vortexing. Fixed cells were incubated at −20 °C for 1 h before pelleting at 400 g for 5 min and washed once in PBS. Mitotic cells were labeled by staining with primary antibody against phospho-Mpm-2 (1:1600) for 1 hr followed by 30 min incubation in secondary antibody conjugated to Alexa Fluor 488 (1:2000). Cells were then centrifuged at 400 g for 5 min and resuspended in PBS with 0.5 μg/ml RNAseA and 10 μg/ml propidium iodide to stain for DNA content. Cell cycle distribution of 10 000 cells was analyzed by using a BD FACSCanto II (BD Biosciences) and BD FACSDiva 9.0.1 software.

### Apoptosis analysis by flow cytometry

For apoptosis analysis, FITC Annexin V Apoptosis Detection Kit with PI (BioLegend) was used according to the manufacturer’s instructions. Briefly, cells were stained with Annexin-FITC and propidium iodide for 15 min at RT after washing the cells once with PBS and then resuspending in Annexin V binding buffer. Dead and Floating cells were added to each sample for analysis by pelleting the conditioned medium at 1500 g for 5 min. Apoptosis of 10 000 cells from each sample was analyzed by using a BD FACSCanto II (BD Biosciences) and BD FACSDiva 9.0.1 software.

### Puromycin assay

To analyze protein translation rate, 16 μg/ml of puromycin were added to siRNA reverse transfected or inhibitor-treated cells as indicated for 45 min (Goodman & Hornberger, 2013). Cell extracts were harvested in RIPA lysis buffer and puromycinylated proteins were detected by western blot using puromycin antibody.

### Statistics

Data were analyzed using GraphPad Prism 6 built-in tests. All data are presented as means±S.D.s. Details about the significance test, the number of replicates, and the *P* values are reported in the respective figure legends.

## Acknowledgments

The authors would like to thank AG Bastians (University of Goettingen) for HeLa cells. We thank Dr. Karen Linnenmannstöns, Dr.Leonie Witte, and Dr. Pradhipa Karuna M for critically reading the manuscript and Mona Honemann-Capito for her excellent technical and scientific support.

## Author contribution

D.C.: Conceptualization, investigation, formal analysis, writing – original draft, review & editing J.C.G.: Conceptualization, Writing – review & editing, funding acquisition

---

**Supplementary Figure 1:**
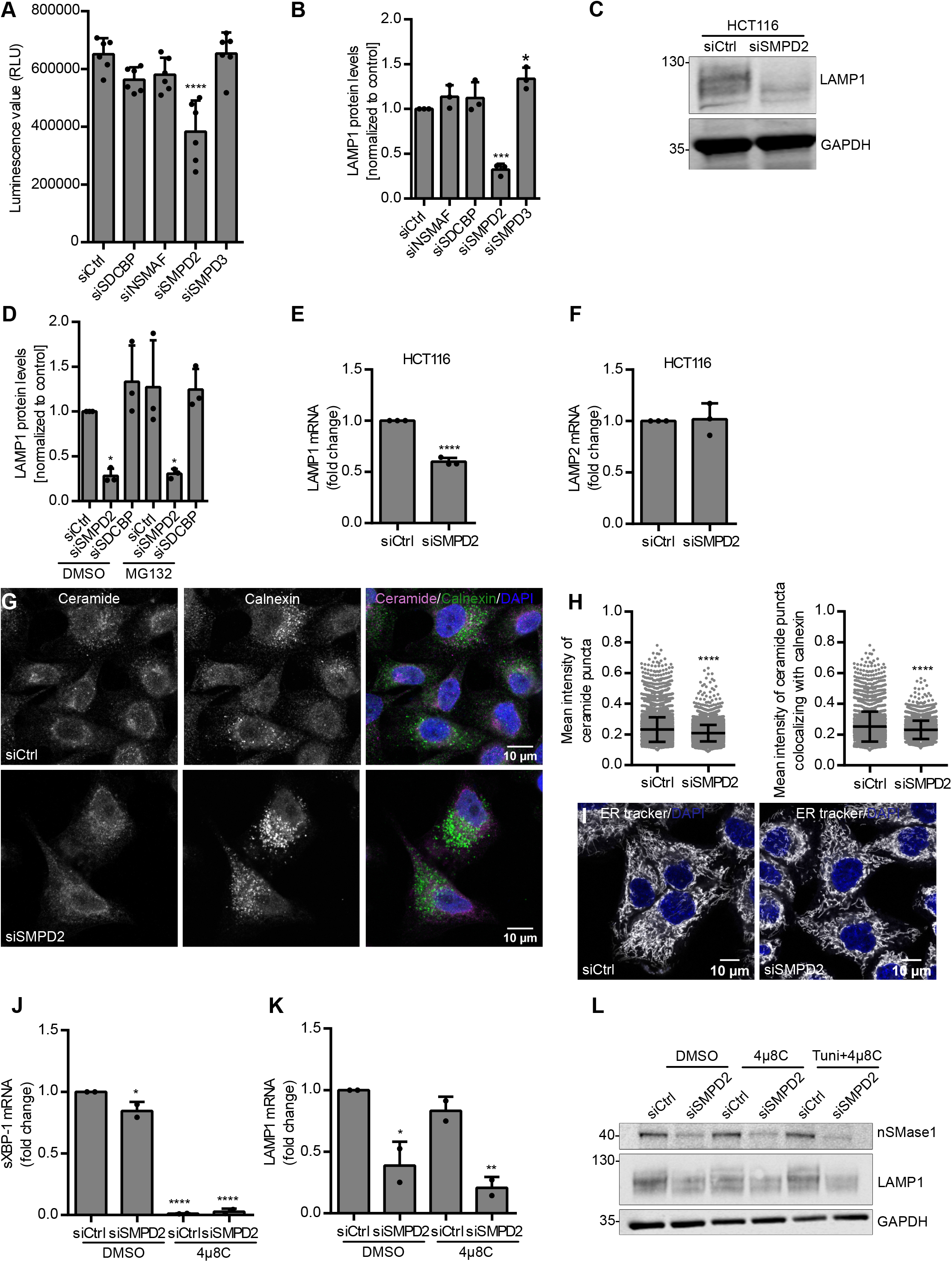
**(A)** Viability assay of HeLa cells transfected with Ctrl, SDCBP, NSMAF, SMPD2, and SMPD3 siRNA for 48 h. Ordinary one-way ANOVA followed by Dunnett’s multiple comparisons test, ****p < 0.0001. **(B)** Western blot quantifications of LAMP1 from (Fig.1A) normalized to the corresponding loading control GAPDH before normalization to the control. Ordinary one-way ANOVA followed by Dunnett’s multiple comparisons test, ***p < 0.0009. **(C)** HCT116 cells were transfected with the indicated siRNAs for 48 h and lysates were immunoblotted against LAMP1 and loading control GAPDH. **(D)** Western blot quantifications of LAMP1 from (Fig.1G) normalized to the corresponding loading control GAPDH before normalization to the control. Ordinary one-way ANOVA followed by Dunnett’s multiple comparisons test, *p < 0.01. (E-F) Real-time qPCR analysis of LAMP1 **(E)** and LAMP2 **(F)** mRNA in HCT116 cells after 72 h of SMPD2 KD. Unpaired Student’s two-tailed *t-test*, ****p < 0.0001. **(G)** HeLa cells transfected with control (Ctrl) and SMPD2 siRNA for 48 h were stained for ceramide and calnexin and analyzed by confocal microscopy. **(H)** Quantifications of mean intensity of ceramide puncta per cell from (G) (left) and mean intensity of ceramide puncta colocalizing with calnexin (right) from three independent experiments. Unpaired Student’s two-tailed *t-test*, ****p < 0.0001. **(I)** HeLa cells transfected with Ctrl or SMPD2 siRNA for 48 h were live imaged by staining with ER tracker and DAPI. **(J)** Real-time qPCR analysis of sXBP-1 mRNA in HeLa cells transfected with Ctrl and SMPD2 siRNA and treated with 4μ8C for 48 h. Ordinary oneway ANOVA followed by Dunnett’s multiple comparisons test, ****p < 0.0001, *p < 0.01, from two independent experiments. **(K)** Real-time qPCR analysis of LAMP1 mRNA in HeLa cells transfected with Ctrl or SMPD2 siRNA and treated with 4μ8C for 48 h. Ordinary one-way ANOVA followed by Dunnett’s multiple comparisons test, ****p < 0.0001, **p < 0.001, *p < 0.01, from two independent experiments. **(L)** HeLa cells reverse transfected with Ctrl and SMPD2 siRNA were grown with DMSO or 4μ8C for 48 h and treated with tunicamycin for 4 h. Cell lysates were immunoblotted against nSMase1, LAMP1, and loading control GAPDH. All experiments from (A)-(L) are done in three biological replicates.

**Supplementary Figure 2:**
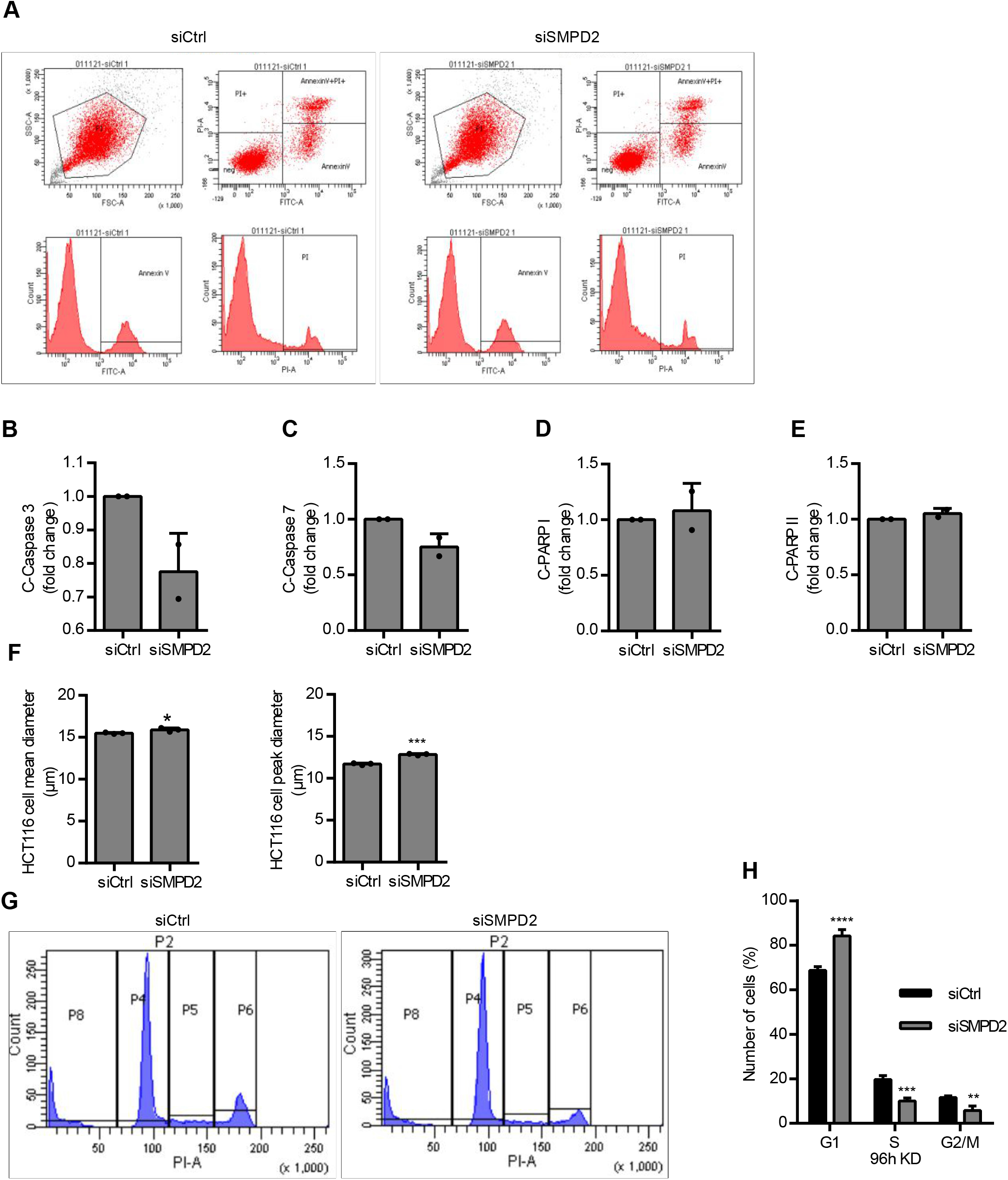
**(A)** Flow Cytometry analysis of FITC Annexin-V and PI staining of HeLa cells transfected with Ctrl (left) or SMPD2 (right) siRNA for 48 h. Forward and side scatter plot (above left) of HeLa cells gated for the subsequent analysis, two-dimensional scatter plot gating cell populations into four quadrants based on PI versus FITC Annexin-V staining (above right), Populations of cells gated for only FITC Annexin-V (below left) and only PI (below right). (B-E) Western blot quantifications of cleaved caspase 3 **(B)**, caspase 7 **(C)**, PARP I **(D)**, and PARP II **(E)** in relative to the respective full length (FL) proteins from two independent experiments. **(F)** HCT116 cell mean diameter (above) and peak diameter (bottom) determined with Casy cell counter after 72 h of Ctrl or SMPD2 siRNA transfection. **(G)** Flow cytometry analysis HeLa cells transfected with Ctrl (left) or SMPD2 (right) siRNA for 48 h were gated to distribute the cell population into G1, S, and G2/M phase of the cell cycle **(H)** Percentage of HeLa cells transfected with Ctrl or SMPD2 siRNA for 96 h were gated for G1, S, and G2/M. Two-way ANOVA followed by Sidak’s multiple comparisons test, ****p < 0.0001, ***p < 0.0009 **p < 0.001. All experiments from (A)-(I) are done in three biological replicates unless otherwise stated.

**Supplementary Figure 3:**
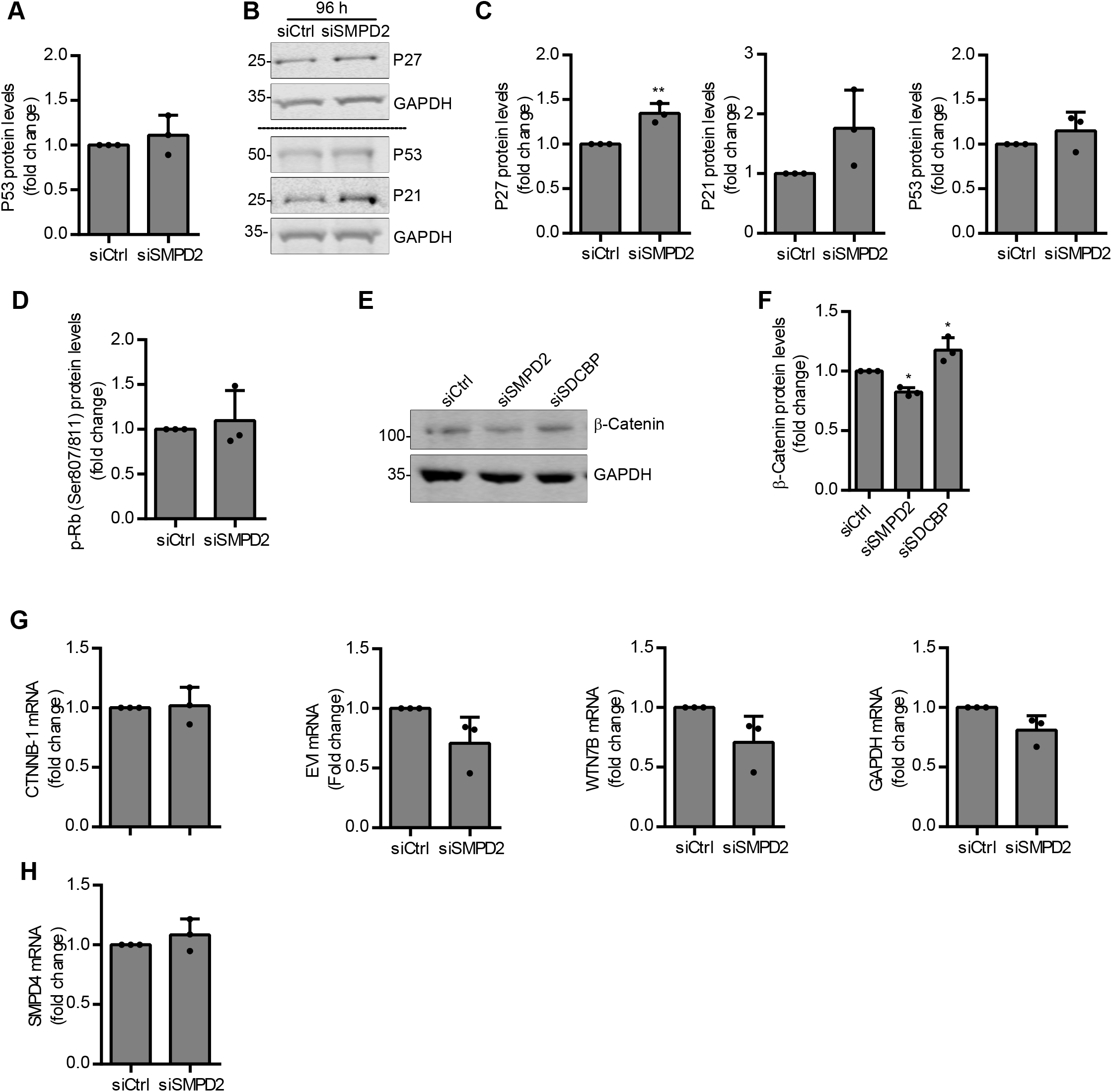
**(A)** Western blot quantifications of P53 protein levels from (Fig. 3G) normalized to the corresponding loading control GAPDH before normalization to the control. Unpaired Student’s two-tailed *t-test*). **(B)** Lysates from HeLa cells transfected with Ctrl or SMPD2 siRNA for 96 h were immunoblotted against P27, P53, P21, and loading control GAPDH. **(C)** Western blot quantifications from (B) of P27, P21, and P53 normalized to the corresponding loading control GAPDH before normalization to the control. Unpaired Student’s two-tailed *t-test*, ****p < 0.0001, **p < 0.001 **(D)** Western blot quantifications of p-Rb from (Fig. 4B) normalized to the corresponding loading control GAPDH before normalization to the control. Unpaired Student’s two-tailed *t-test*, **p < 0.001. **(E)** HeLa cells were transfected with the indicated siRNAs for 48 h and lysates were immunoblotted against β-catenin and loading control GAPDH. **(F)** Western blot quantifications of β-catenin from (E) normalized to the corresponding loading control GAPDH before normalization to the control. Ordinary one-way ANOVA followed by Dunnett’s multiple comparisons test, *p < 0.01. **(G)** Real-time qPCR analysis of CTNNB-1, EVI, WNT7B, and GAPDH mRNA in HCT116 cells transfected with Ctrl and SMPD2 siRNA for 72 h. Unpaired Student’s two-tailed *t-test*. (**H)** Real-time qPCR analysis of SMPD4 mRNA in HCT116 cells transfected with Ctrl and SMPD2 siRNA for 72 h. Unpaired Student’s twotailed *t-test*, ****p < 0.0001. All experiments from (A)-(H) are done in three biological replicates.

